# Novel Approach for Parallelizing Pairwise Comparison Problems as Applied to Detecting Segments Identical By Decent in Whole-Genome Data

**DOI:** 10.1101/2020.07.07.191999

**Authors:** Emmanuel Sapin, Matthew C. Keller

## Abstract

**Motivation:** Pairwise comparison problems arise in many areas of science. In genomics, datasets are already large and getting larger, and so operations that require pairwise comparisons—either on pairs of SNPs or pairs of individuals—are extremely computationally challenging. We propose a generic algorithm for addressing pairwise comparison problems that breaks a large problem (of order *n*^2^ comparisons) into multiple smaller ones (each of order *n* comparisons), allowing for massive parallelization.

**Results:** We demonstrated that this procedure is very efficient for calling identical by descent (IBD) segments between all pairs of individuals in the UK Biobank dataset, with a user time savings roughly 180-fold over the traditional (non-parallel) approach to detecting such segments. This efficiency should extend to other methods of IBD calling and, more generally, to other pairwise comparison tasks in genomics or other areas of science.

## 1 Introduction

Situations where all pairs of observations in a dataset must be compared arise in many areas of science. For example, in protein studies, forming the graphs used in protein clustering relies on finding a protein’s likeness to every other protein (Chapman and Kalyanaraman 2011; Sapin *et al*. 2016), and in physics, calculating the total force each body has on every other body is required in order to predict the position and motion of all bodies in the n-body problem (Leimanis and Minorsky 1958). Such pairwise comparison problems are extremely computationally challenging with large datasets because they grow at the square of the sample size (are of order 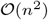).

Because genomics datasets are growing quickly in both numbers of samples and numbers of measured variants, the computational challenge inherent in comparing all pairs of observations is particularly acute in several types of genomic analyses, such as the detection of epistasis (Sapin *et al*. 2014; Fang *et al*. 2012) and the construction of gene regulatory networks (Chang *et al*. 2008; Qiu *et al*. 2009). Another such example in modern genomics, and the motivation for the procedure introduced in this manuscript, is the detection of identical by descent (IBD) segments between all pairs of individuals using whole-genome single nucleotide polymorphism (SNP) data (Browning and Browning 2012). Two haplotypes (homologous chromosomal segments of DNA) are IBD if they descend from a common ancestor without either haplotype experiencing an intervening recombination (Powell *et al*. 2010). IBD segments can be used for a number of downstream analyses in genetics, including imputation, phasing, inference of the degree of relatedness, IBD mapping to detect the effects of rare variants on phenotypes, estimation of the effective population sizes, and detection of signatures of recent positive selection (Bjelland *et al*. 2007). IBD detection requires that each pair of individuals in a dataset is compared each location across the genome, typically in phased data (where the homologous chromosomes inherited from the father and mother have been computationally distinguished from each other). Segments that match for a stretch that is too long to have arisen by chance are deemed “IBD.” This procedure is usually parallelized by splitting the genome into 23 subsets corresponding to the 23 chromosomes, but further subsetting by smaller genomic (sub-chromosomal) windows is problematic because it increases the miss rate of IBD segments that span two or more windows, because it adds a computationally expensive post-processing step (stitching together IBD segments that span across windows) (Browning and Browning 2012), and because each window still requires 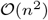 comparisons, which can become too computationally expensive in large datasets even if the windows are small.

Several methods have been proposed to compare all pairs in a dataset in efficient ways, although none of these have been applied to IBD detection to our knowledge. While some are applicable only to specific problems (Parikshit *et al*. 2009; Cormen *et al*. 1990; Wauthier *et al*. 2013; Krejčí 2018), a general strategy that can be applied across multiple applications is to parallelize the comparisons, hopefully in some kind of efficient manner. Kiefer *et al*. (Kiefer *et al*. 2010) reviewed three ways to accomplish this. The first (“broadcast”) approach consists solely in having multiple processes performing comparisons each on a subset of pairs. The second (“block”) approach subsets pairs that are more likely to have elements in common, which limits data replication (see also (Kleinheksel and Somani 2016)). The final (“design”) approach uses a projective plane (Lee *et al*. 2004) or *symmetric balanced incomplete block design* to create subsets of elements such that any pair of elements is observed exactly once across all the subsets.

In the current manuscript, we introduce a procedure for parallelizing the pairwise comparison problem, similarly to the “design” approach in (Kiefer *et al*. 2010), by subsetting the set of individuals such that each pair is observed exactly once across all the subsets. Here an affine plane (Hughes and Piper 1973) is used for this purpose, creating slightly more subsets 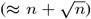 than the *n* individuals in the set. The sample size for each subset 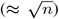 is roughly midway between the naive approach (2 individuals in each of *n*(*n* + 1)/2 subsets) and the brute force approach (a single set of *n* individuals entailing *n*(*n* + 1)/2 comparisons, including self-comparisons), which is an excellent middle ground for massive and efficient parallelization for many applications.

Applying our approach to the detection of IBD segments on genome-wide SNP data from 435,187 individuals from the UK Biobank dataset (Bycroft *et al*. 2018), we show that our procedure resulted in a large saving in time over alternative approaches. Our procedure should allow for efficient parallelization of IBD detection in much larger datasets than this one, and the general approach is easily extendable to other applications outside of IBD detection and genomics.

## 2 Parallelization Methodology

We describe here how to create subsets for samples of any size made up of individuals, *x_i_, i* ∈ {1…*n*} with *n*>=4, such that all pairs of individuals, {*x_i_, x_j_*}, are observed exactly once across all subsets. Let *p* be the lowest prime number such that *p*^2^ ≥ *n*. To create the subsets, the numbers from 1 to *p*^2^ are placed column-wise in a *p*-by-*p* matrix *P* as shown in Figure **??**. Numbers 1…*n* index an individual in the sample, and numbers *n* + 1…*p*^2^, if any, are unassociated (null), which can lead to some subsets having fewer individuals than others.

Our procedure leads to a total of 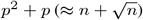 unique subsets of individuals. The first *p* subsets are defined by each of the columns of matrix *P* and the next *p* subsets are defined by each of the rows of matrix *P*, for a total of 2*p* subsets drawn from matrix *P*. For example, the final subset based on the rows of *P* contains individuals *x_p_, x*_2*p*_, *x*_3*p*_, …*x*_*p*^2^_.

The *p*^2^ + *p* − 2*p = p*^2^ − *p* remaining subsets are defined by new matrices, *P_k_*. Each matrix *P_k_* (*k* ∈ {1…*p* − 1}) is created via shifting operations on the elements in the columns of *P*, as described below. Unlike the original 2*p* subsets based on *P*, subsets based on *P_k_* are defined only by each of the rows (not columns) of *P_k_*, and thus each *P_k_* matrix contributes *p* additional subsets. To create the *p* − 1 *P_k_* matrices, the elements of each column *i* of *P_k_* are shifted up relative to the elements in the corresponding column *i* of *P* by *r*(*i,k*) rows, where

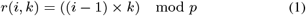

and mod is the modulo operation. For example, for *k* =1, *r*(*i* = 1, *k* = 1) = 0 and so the first column of *P*_1_ is identical to the first column of *P, r*(*i* = 2, *k* = 1) = 1 and so the second column of *P*_1_ is shifted by one element relative to column 2 of *P*, and *r*(*i* = 3, *k* = 1) = 2 and so the third column of *P*_1_ is shifted by two elements relative to column 3 of *P*, and so forth. This shifting is defined such that the elements that were at the top of column *i* in *P* are shifted to the bottom of column *i* in *P_k_*, as shown Figure 1. For example, the last element of the second column of *P*_1_ is the first element of the second column of *P* (*P*_1_ [*p*, 2] = *P*[1, 2]), and the last two elements of the third column of *P*_1_ are the first two elements of the third column of *P*. Thus, the total number of subsets is *p*^2^ + *p* (of which 2*p* are created from the original matrix *P* and the remainder are created from shifted matrices *P_k_*), each individual is in *p* + 1 different subsets, and the number of individuals per subset is *p* (or fewer for some subsets when *p*^2^ > *n*). Figure 2 shows the matrices *P, P*_1_, *P*_2_, *P*_3_, and *P*_4_ for an example where *n = p*^2^ = 25.

**Figure 1:**
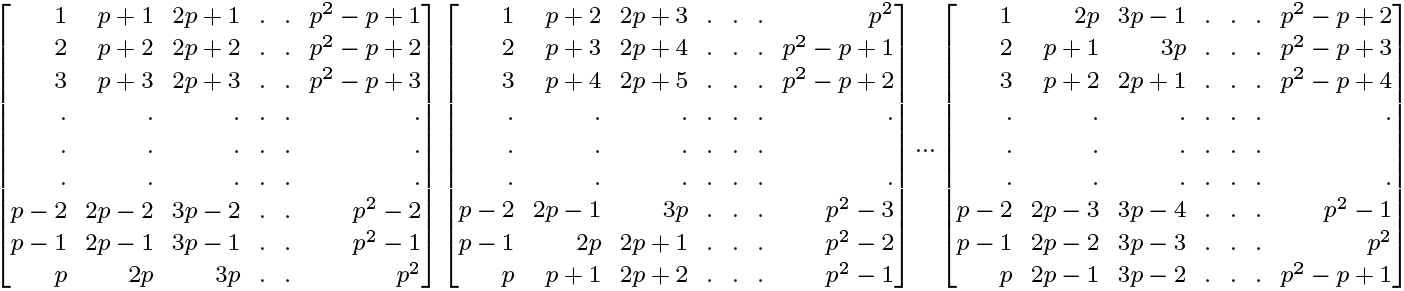
Matrix *P*, *P*_1_, and *P*_*p*−1_

**Figure 2:**
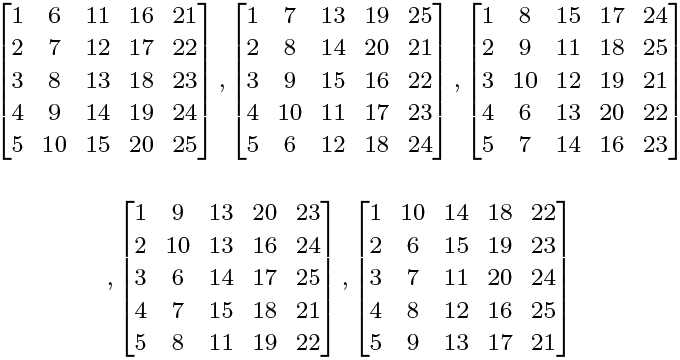
Matrices *P*, *P*_1_, *P*_2_, *P*_3_, and *P*_4_ for *n* = *p*^2^ =25

To assess this procedure works as intended across a range of *n*, we counted the number of times *t* each pair of individuals was in the same subset across all subsets. As any pair should be in one and only one subset, *t* should be equal to 1 for all pairs, and this is what we observed for all *n* < 500*K*. Although we have found no formal proof that this algorithm works to always define *p*^2^ + *p* subsets such that each pair of individuals is in exactly one subset, we have computationally demonstrated that the algorithm works in this way for any *n* < 500*K* and have no reason to suspect it would behave differently for *n* > 500*K*. Therefore the collection of these subsets forms an affine plan where subsets are lines and individuals in a subset are points on the line. A GitHub page is available at https://github.com/emmanuelsapin/Subsets-for-the-Parallelization-of-All-Pair-Comparaison-Algorithms with the code to generate lists of IDs of individuals composing each subset.

The total number of comparisons (including self-comparisons) made under our approach is 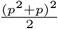, the product of the *p*^2^ + *p* subsets and the 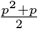 comparisons per subset. The total number of comparisons under the naive approach, where all pairs are compared in the entire dataset, is 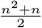. Thus, under the optimal scenario, where *p*^2^ = *n* exactly, our approach requires only a total of 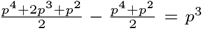 (or 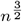) more comparisons than the naive approach. When *p*^2^ > *n* and *n* is large (e.g., > 100K), the number of comparisons made by our approach is proportionately only slightly greater than this. Thus, our algorithm requires roughly the same number of pairwise comparison computations as the naive approach, but with the benefit of massive parallelization.

Algorithm 1 describes the full process.

**Algorithm 1:**
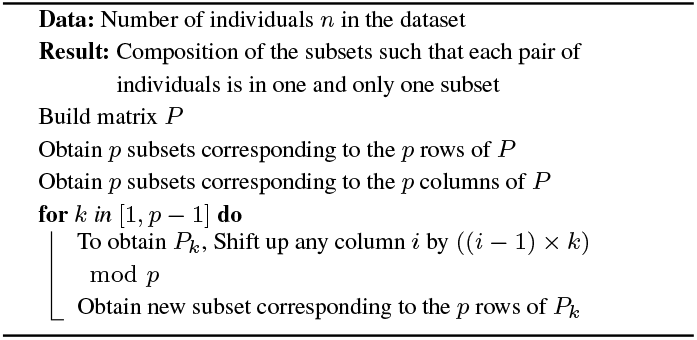
Main Algorithm

## 3 Implementation and results

We used our procedure to parallelize the detection of IBD segments in a whole-genome (*m* = 330,005 SNPs) dataset of 435,187 individuals from the UK Biobank project (Bycroft *et al*. 2018). For this sample size, the lowest prime number, *p*, such that *p*^2^ ≥ 435, 187 was 661, and thus *p* = 661 resulting in 661^2^ + 661 = 437, 582 subsets. We called IBD segments > 3.5cM (centi-Morgans) in length using the GERMLINE software (Gusev *et al*. 2009), which is among the most efficient and accurate estimators of IBD (Bjelland *et al*. 2007). We ran 1,000 instances of GERMLINE in parallel for each of 438 total jobs. In addition to the parallelization, we modified GERMLINE to utilize shared memory segments across tasks for each job, such that the genomic data was read from the hard drive only once per job. These jobs were run on 140 nodes of the Blanca Condo Cluster at the University of Colorado at Boulder (https://www.colorado.edu/rc/res-ources/blanca). The total CPU hour usage for this task was 5,644 hours, which is higher than the CPU hour usage would have been had we used GERMLINE on the full sample, subsetting only across chromosomes (estimated to be 4,272 CPU hours). However, massively parallelizing the job translated into a very large savings in user hours. The final job ended 28.5 hours after the first one was submitted. 34 jobs failed for technical reasons (e.g., lack of space on the local hard drive) and were re-run the following days. Thus, about 99.64% of the task was achieved in under 29 hours.

We estimate that running GERMLINE in parallel only across the 23 chromosomes would have required more than a terabyte of memory and over six months to complete using a 2x Intel Xeon Gold 6130 (2.1 GHZ) processor under a x86_64 architecture, whereas running GERMLINE in parallel on every pair independently would have taken even longer due to the startup cost for each of ≈ 9.5 × 10^10^ total processes. Using our novel procedure, we achieved this task in under 29 hours. The IBD segments we obtained will be used to infer the degree of relatedness between all pairs of individuals in our dataset, which is another pairwise comparison problem that can be optimized using the same procedure introduced here.

## 4 Conclusion

We developed a parallelisation strategy that subsets individuals in order to break a large pairwise comparison problem (of order *n*^2^ comparisons) into multiple smaller pairwise comparisons problems (each of order *n* comparisons).

Each pair of individuals is compared exactly once across all subsets and the total number of comparisons that must be made under our algorithm is only slightly higher than the number that must be made if all pairs were compared in the entire dataset without subsetting.

We demonstrated that this procedure is very efficient for calling IBD segments using GERMLINE in the large UK Biobank dataset, with a user time savings roughly 180-fold over running GERMLINE on the entire sample in our particular instance. The amount of time saved using our procedure will depend on the particular algorithms being used and on the computational architecture, but to the degree that the algorithms scale over order *n*, and especially as they approach order *n*^2^, as many algorithms that work at the unit of pairs should, our procedure should offer substantial savings in user-time. While our procedure was highly efficient for calling IBD segments using GERMLINE, there are good reasons to believe that this efficiency will extend to other methods of IBD calling and, more generally, to other pairwise comparison tasks in genomics or other areas of science.

## 6 Acknowledgment

This research was conducted using the UK Biobank Resource under application numbers 16651. This work utilized resources from the University of Colorado Boulder Research Computing Group, which is supported by the National Science Foundation (awards ACI-1532235 and ACI-1532236), the University of Colorado Boulder, and Colorado State University. We thank Mr. Jared Balbona and Drs. Luke Evans and Richard Border for their extensive help throughout this project.

